# The role of hippocampal spatial representations in contextualization and generalization of fear

**DOI:** 10.1101/514505

**Authors:** Lycia D. de Voogd, Yannick P. J. Murray, Ramona M. Barte, Anouk van der Heide, Guillén Fernández, Christian F. Doeller, Erno J. Hermans

## Abstract

Using contextual information to predict aversive events is a critical ability that protects from generalizing fear responses to safe contexts. Animal models have demonstrated the importance of spatial context representations within the hippocampal formation in contextualization of fear learning. The ventromedial prefrontal cortex (vmPFC) is known to play an important role in safety learning, possibly also through the incorporation of context information. However, if contextual representations are related to context-dependent expression of fear memory in humans remains unclear. Twenty-one healthy participants underwent functional MRI combined with a cue-context conditioning paradigm within a self-navigated virtual reality environment. The environment included two buildings (Threat and Safe context), which had distinct features outside but were identical inside. Within each context, participants saw two cues (CS+, CS−). The CS+ was consistently (100% reinforcement rate) paired with an electric shock in the Threat context, but never in the Safe context. The CS− was never paired with a shock. We found robust differential skin conductance responses (SCRs; CS+ > CS−) in the Threat context, but also within the Safe context, indicating fear generalization. Within the Safe context, vmPFC responses to the CS+ were larger than those in the Threat context. We furthermore found environment-specific representations for the two contexts in the training paradigm (*i.e*., before conditioning took place) in the hippocampus to be related to fear expression and generalization. Namely, participants with a weak context representation (z-score < 1.65) showed stronger fear generalization compared to participants with a strong context representation (z-score > 1.65). Thus, a low neural representation of spatial context may explain overgeneralization of memory to safe contexts. In addition, our findings demonstrate that context-dependent regulation of fear expression engages ventromedial prefrontal pathways suggesting this involves a similar mechanism that is known to be involved in retrieval of extinction memory.

## Introduction

The unwanted expression of learned fear responses in safe environments is one of the hallmarks of anxiety and fear-related disorders (Lissek & Grillon, 2010; Shin & Liberzon, 2010). A crucial factor that protects against this overgeneralization of fear is the ability to correctly use contextual factors to predict threat and regulate fear expression. Animal research has shown that accurate context-dependent prediction of threat depends on spatial context representations within the hippocampal formation (Maren, Phan, & Liberzon, 2013). The ventromedial prefrontal cortex (vmPFC) is known to be critically involved in regulating fear expression (Dunsmoor & Paz, 2015; Greenberg, Carlson, Cha, Hajcak, & Mujica-Parodi, 2013), but also fear generalization (Xu & Südhof, 2013). In humans, the role of these regions in context-dependent expression and generalization of fear remains to be investigated.

The role of context information in fear learning has been mostly studied in animal models using context conditioning paradigms (Maren et al., 2013). In these paradigms, rodents learn that the context (*e.g*., a box) predicts an aversive outcome, unlike cue conditioning paradigms, in which a cue (*e.g*., a tone) serves as conditioned stimulus (CS) to predict an unconditioned stimulus (UCS). Different neural mechanisms are involved in associating threat with a cue versus a context. The amygdala plays a critical role in both the formation and expression of the cue CS-UCS association (LeDoux, 2003; Phillips & LeDoux, 1992), although in humans, the role of the amygdala in fear expression is less clear (Fullana et al., 2016; Mechias, Etkin, & Kalisch, 2010). The hippocampus is critically involved in spatial mapping (Epstein, Patai, Julian, & Spiers, 2017; O’Keefe & Nadel, 1978) and thereby also plays a role in the acquisition and storage of contextual threat information (Wiltgen, 2006). In addition to the context predicting the aversive outcome by itself, the context can also change the predictive value of the cue. In this case, the context serves as an “occasion setter” (Bouton & Nelson, 1998; Holland & Bouton, 1999; Maren et al., 2013) such that the CS predicts an aversive outcome in one context, but not in another.

Electrophysiological studies in rodents have demonstrated the existence of *place* cells within the hippocampus which selectively increase their firing rates when an animal is at a specific location (O’Keefe & Dostrovsky, 1971), thus providing a neuronal representation of the spatial environment (O’Keefe & Conway, 1978; Wilson & McNaughton, 1993). In addition to neuronal spikes, hippocampal local field potentials (LFP) have also been shown to code spatial location (Agarwal et al., 2014). Similarly, research with intracranial recordings in humans has shown that cells in the hippocampus respond to specific spatial locations while navigating around a virtual environment (Ekstrom et al., 2003). Also using non-invasive imaging techniques, such as blood oxygen level-dependent functional magnetic resonance imaging (BOLD-fMRI), evidence was found for spatial representations in medial temporal lobe (MTL) regions in humans (Steemers et al., 2016). Work using machine learning methods applied to BOLD-fMRI data has furthermore shown that it is possible to decode environment-specific representantions within a virtual environment from patterns of hippocampal activity (Hassabis et al., 2009; Rodriguez, 2010; Sulpizio, Committeri, & Galati, 2014). We therefore reasoned that the ability to decode environment-specific representantions from such patterns of hippocampal activity can serve as an index of the strength of a context representation.

Context representations might play an important role in fear generalization. Animal research has shown that lesions in the hippocampus prevent contextual fear conditioning in rats (Kim & Fanselow, 1992), providing direct evidence for the importance of the hippocampus for contextual fear learning. In humans, using virtual reality on a computer screen (Dunsmoor, Ahs, Zielinski, & LaBar, 2014) or a head-mounted device (Kroes, Dunsmoor, Mackey, McClay, & Phelps, 2017), it is now possible to study the role of context in fear learning. Initial studies using virtual reality by means of preprogrammed movies have shown that responses to the CS+ acquired in a threat context generalize to a safe context (Baas, Nugent, Lissek, Pine, & Grillon, 2004). However, since volitional movement influences the formation of context representations in rodents (Cei, Girardeau, Drieu, Kanbi, & Zugaro, 2014), these paradigms are less suited to investigate context representations. Here, we therefore used a self-navigated virtual reality environment to test the hypothesis that individuals with a low hippocampal context representation show increased fear generalization in the safe context compared to those with a high context representation.

The vmPFC is thought to play a critical role in regulating the expression of fear memory, and has been investigated extensively in the context of extinction leaning (Milad & Quirk, 2012). It has been shown that extinction recall is context dependent (Milad, Orr, Pitman, & Rauch, 2005), and that this context-dependent expression of extinction requires the vmPFC and its interactions with the hippocampus and amygdala (Maren et al., 2013). A failure to recruit the vmPFC in response to safe stimuli has been associated with deficiencies in fear generalization (Cha et al., 2014; Holt, Coombs, Zeidan, Goff, & Milad, 2012). In animal models, the mPFC has been implicated in fear generalization as well (Xu & Südhof, 2013). We reasoned that the presentation of a CS+ in a safe context could lead to a suppression of fear responses similar to a CS+ presentation during extinction recall (Maren et al., 2013) and therefore hypothesized that processing of a cue predicting an aversive outcome presented in a safe context is supported by similar vmPFC-dependent mechanisms as those involved in extinction recall.

The present study was designed to test these hypotheses in humans using fMRI and a virtual reality environment. Twenty-one healthy participants underwent a context training paradigm within the virtual reality environment. This environment involved a rich landscape including two buildings. The buildings had distinct features on the outside, but were identical on the inside. Within this environment, participants underwent context training which was followed by a differential delay contextual fear conditioning paradigm with spatial context as occasion setter. The context training paradigm served to create a strong context representation, and to allow for measurement of these representations without the potential confound of threat-induced arousal during the fear conditioning paradigm. During the fear conditioning paradigm, two different cues (CS+ and CS−) were presented in both buildings, but the CS+ was only paired with a mild electric shock (at a reinforcement rate of 100%) in the threat context, and never in the safe context. The CS− was never paired with a shock. Estimations of spatial context representation strength were obtained using a linear support vector machine (SVM) classifier that we trained on hippocampal patterns obtained from and tested in the context training paradigm.

We tested the following predictions. First, we expected that, in addition to differential conditioned SCRs in the threat building, differential conditioned SCRs in the safe building would be present (i.e., where participants never received a shock, indicating fear generalization). Second, we expected that also differential conditioned BOLD responses in the threat context would generalize to the safe context. Third, if context-dependent downregulation of fear expression in safe environments is indeed a process similar to extinction recall, then CS+ in the safe context should elicit stronger BOLD responses in the vmPFC than CS+ in the threat context. Fourth, we predicted using multi-voxel patterns of hippocampal activity that the hippocampus has a distinct representation of the two buildings in the virtual environment. If low representations of spatial context play a role in fear overgeneralization, then participants with lower classifier accuracy in distinguishing the two buildings should exhibit increased generalization of conditioned SCRs the safe context.

## Methods

### Participants

Twenty-one right-handed healthy volunteers (12 females, 12 males; 19-34 years [M=23.3, SD=3.6]) completed the study. Exclusion criteria were: current or lifetime history of psychiatric, neurological, or endocrine illness, current treatment with any medication that affects central nervous system or endocrine systems, average use of more than three alcoholic beverages daily, average use of recreational drugs weekly or more, habitual smoking, predominant left-handedness, uncorrected vision, intense daily physical exercise, and any contraindications for MRI. Participants gave written informed consent and were paid for their participation. This study was approved by the local ethical review board (CMO region Arnhem-Nijmegen).

### Design and procedure

The design of this study is illustrated in ***Figure 1***. Each session started with a familiarization of the virtual reality task outside of the scanner. This was done to minimize learning and/or novelty effects during the scanning phase and to allow time for context representations to be formed and stabilized, and create a “cognitive map” (Doeller, King, & Burgess, 2008; Hassabis et al., 2009). Next, participants underwent a functional MRI session which included the following tasks: an object localizer task (7 min 40 sec), the context training paradigm (16-21 min), and the fear conditioning paradigm (25-30 min). The localizer paradigm was programmed using Presentation^®^ software (Version 0.70, www.neurobs.com). Data including the localizer paradigm will be reported elsewhere. The virtual reality environment was built and presented using the Unreal Development Kit (Unreal Engine 3, http://udn.epicgames.com/Three/WebHome.html) which communicated with the Presentation^®^ software to be able to synchronize with the image acquisition and trigger the electrical shocks in the fear conditioning paradigm.

**Figure 1.**
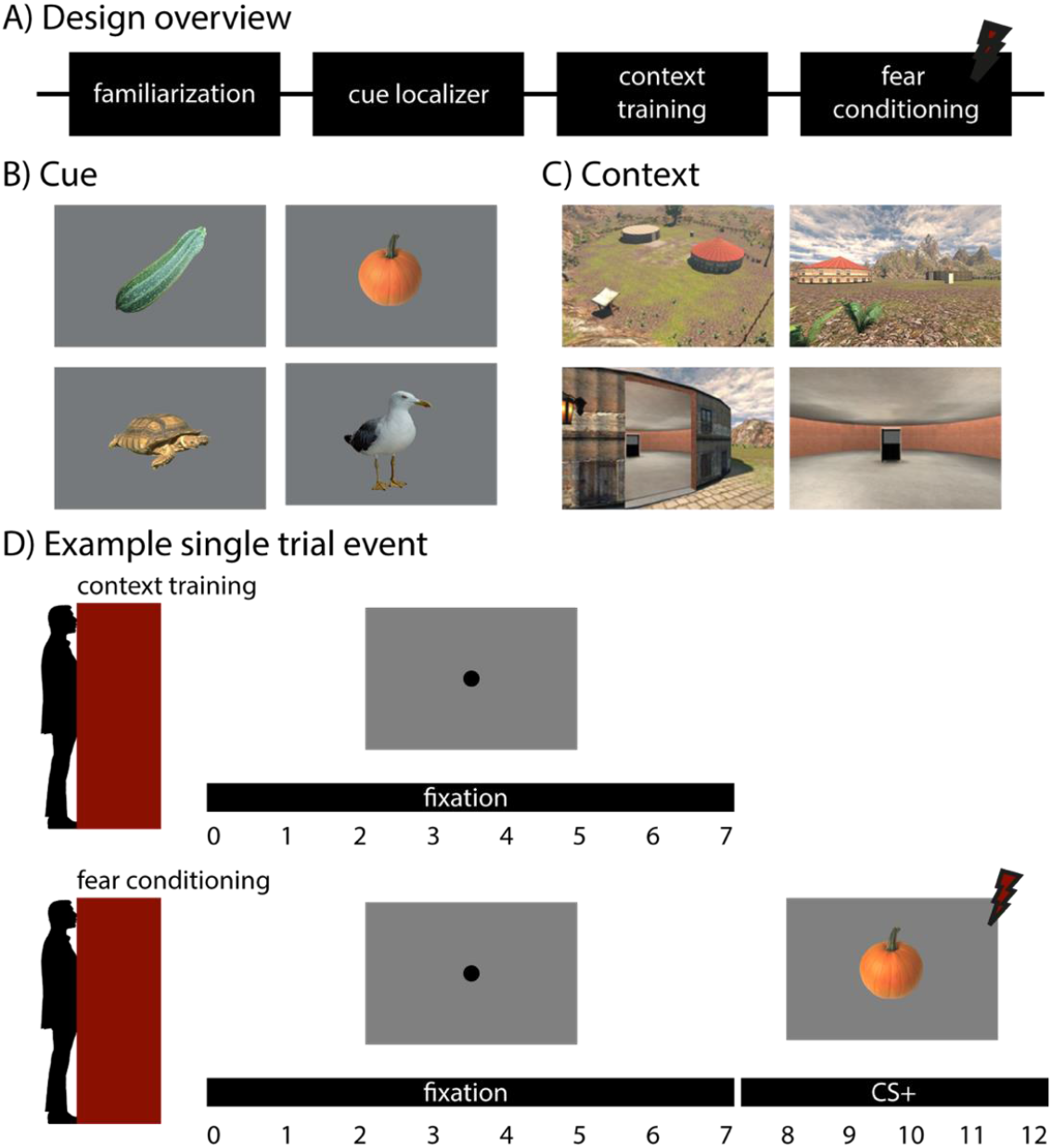
Overview of the experimental design. (A) The task consisted of four stages. First, participants were familiarized with the virtual reality environment outside the MRI. Next, participants underwent a functional MRI session in which they performed an object localizer paradigm including different versions of the CSs, a context training phase in the virtual environment, and finally a fear conditioning phase. Both phases included two runs. (B) Illustration of the conditioned stimuli. Each participant saw either a zucchini or a pumpkin (CS+ or CS− counterbalanced) and either a turtle or seagull (CS+ or CS− counterbalanced). (C) An overview of the virtual reality environment. Participants navigated through it from a first-person perspective. The environment contained two buildings (Threat or Safe counterbalanced). (D) A schematic overview of a trial within a building for the context training phase and the fear conditioning phase. When participants entered the “Trial booth”, depicted as a red rectangle, within the building, the 3D environment converted to a 2D environment. A fixation dot appeared on the screen for seven seconds. In the fear conditioning phase, this was followed by the CS for five seconds. In the threat building, the CS+ was always followed by a shock.

### Context familiarization of the virtual environment

Participants navigated through a virtual environment from a first-person perspective. The environment involved a rich landscape including two buildings which they were able to enter. See ***Figure 1C*** for an illustration of the environment. The buildings had distinct features on the outside but were identical on the inside. First, we familiarized participants with the entire virtual environment which was done outside the MRI scanner. In this paradigm, participants navigated through the environment using three buttons (*i.e*., forward, left, and right) based on instructions that where given to them. For example, participants were instructed to “go to the wooden barrel” or “walk to the bucket”. Through these instructions, participants visited all areas of the environment at least once.

After this initial familiarization paradigm, the participants practiced the main task. They were placed in the environment and were instructed to walk to an instruction booth located outside of the two main buildings. Once they entered, the number “1” or “2” was displayed, indicating which building they were required to visit. Participants had to figure out which building corresponded to this number. If they stood in front of the door of the correct building, the door would open and they had to enter. In the middle of the building, there was a “trial booth” which they had to enter. At that moment, participants could not navigate anymore and a trial would be presented on the screen in 2D. For this familiarization paradigm, a trial consisted of a fixation cross presented for 4-6 sec. In total, there were 48 trials (24 per building). If after a trial, the next trial was in the same building, the door would stay closed. There were always two to four consecutive trials in the same building to reduce the amount of time spent on navigation. After these trials, the door would be open and participants could leave the building and navigate to the instruction booth, which switched to a new random location each time, but was always outside the two buildings. Participants never visited the same building more than twice in a row.

### Context training of the virtual environment during functional MRI

Once in the MRI scanner, participants were placed in the virtual environment again. This paradigm was very similar to the last part of the familiarization paradigm. The instruction booth indicated which building they had to enter. Once the participant entered the building they were instructed to go to the trial booth in the middle of the building. Once they entered the booth, a trial started. In this paradigm, a trial consisted only of a gray screen with a fixation cross presented for 7 s. This was done to sample hippocampal response patterns within the two buildings in the absence of a stimulus, and with identical visual features at the moment of sampling. After the end of a trial there were again two options, either to go to the booth in the same building again, or to go to the instruction booth. There were always two to four consecutive trials within the same building, no more than twice they had to go to the same building, and in total there were 48 trials (24 per building) divided over two scan runs. Navigating through the environment was self-paced, therefore the duration of the task varied between participants (range: 16-21 min, M=18 min 10 sec, SD=1 min 30 sec).

### Fear acquisition in the virtual environment during functional MRI

The fear conditioning paradigm took place in the same environment and was therefore very similar in setup to the context training paradigm. Here, a trial consisted of a fixation cross with a duration of 7 s, which was followed by a conditioned stimulus (CS) presentation with a duration of 5 s. In each trial, one of two CSs, a CS+ and a CS−, was shown. The same CSs were used in each of the two buildings, but only in one of the buildings (the Threat context), the CS+ predicted an electrical shock. In the other building (the Safe context) participants never received a shock. The CS− was never paired with shock. There were two pairs of CS+/CS− (a picture of a Seagull/Pumpkin or Tortoise/Zucchini) the use of which was counterbalanced across participants. See ***Figure 1B*** for an illustration. Again, after each trial, there were two options, either to stay in the same building for a new trial, or to leave the building and go to the instruction booth. There were always two to four consecutive trials within the same building, no more than twice they had to go to the same building, and in total there were 60 trials (30 per building), divided over two scan runs. Because navigation through the environment was self-paced, the duration of this paradigm varied between participants (range: 25-30 min, M=27 min 30 sec, SD=1 min).

### Peripheral stimulation

Electrical shocks were delivered via two Ag/AgCl electrodes attached to the distal phalanges of the thumb and fifth digit of the right hand using a MAXTENS 2000 (Bio-Protech) device. Shock duration was 200 ms, and intensity varied in 10 intensity steps between 0V-40V/0mA-80mA. During a standardized shock intensity adjustment procedure, each participant received and subjectively rated five shocks, allowing shock intensity to converge to a level experienced as uncomfortable, but not painful. The resulting average intensity step was 4.1 (SD: 1.5) on a scale from 1 to 10.

### Peripheral measurements

Electrodermal activity (EDA) was assessed using two Ag/AgCl electrodes attached to the distal phalanges of the second and third digit of the left hand using a BrainAmp MR system and recorded using BrainVision Recorder software (Brain Products GmbH, Munich, Germany). Data were preprocessed using in-house software; radio frequency (RF) artifacts were removed and a low-pass filter was applied (de Voogd, Fernández, & Hermans, 2016b, 2016a). Skin conductance responses (SCR) were automatically scored with additional manual supervision using Autonomate (Green, Kragel, Fecteau, & LaBar, 2014) implemented in Matlab 7.14 (MathWorks). SCR amplitudes (measured in μSiem) were determined for each trial within an onset latency window between 0.5 and 3.8 s after stimulus onset, with a minimum rise time of 0.5 s and because there was a 100% reinforcement rate, the maximum latency of the peak was set at 4.9 s, the time point the shock was given. All response amplitudes were square-root transformed and normalized according to each participant’s mean UCS response (Schiller, Kanen, LeDoux, Monfils, & Phelps, 2013) prior to statistical analysis. Analyses on the SCRs were performed using SPSS 23 (IBM Corp, Armonk, New York). A repeated-measures ANOVA was conducted including CS (CS+, CS−), Context (Threat, Safe) as within-subject factors. To assess fear generalization, we calculated a fear-generalization index by subtracting the differential SCR responses from the last two trials in the safe context from the differential SCR responses from the last two trials in the threat context. Effect sizes are reported as Partial eta squared 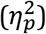 and Cohen’s D.

Finger pulse was recorded using a pulse oximeter affixed to the second digit of the left hand. Respiration was measured using a respiration belt placed around the participant’s abdomen. Pulse and respiration measures were used for retrospective image-based correction (RETROICOR) of physiological noise artifacts in BOLD-fMRI data (Glover, Li, & Ress, 2000). Raw pulse and respiratory data were processed offline using in-house software for interactive visual artifact correction and peak detection, and were used to specify fifth-order Fourier models of the cardiac and respiratory phase-related modulation of the BOLD signal (Van Buuren et al., 2009), yielding 10 nuisance regressors for cardiac noise and 10 for respiratory noise. Additional regressors were calculated for heart rate frequency, heart rate variability, (raw) abdominal circumference, respiratory frequency, respiratory amplitude, and respiration volume per unit time (Birn, Diamond, Smith, & Bandettini, 2006), yielding a total of 26 nuisance regressors.

### MRI data acquisition

MRI scans were acquired using a Siemens (Erlangen, Germany) MAGNETOM Skyra 3.0T MR scanner with a 32-channel head coil. T2*-weighted BOLD images were recorded using a multiband accelerated Echo-planar imaging (EPI) sequence [TR, 0.909 s; TE, 24.6 ms; Generalized Autocalibrating Partially Parallel Acquisitions (GRAPPA; Griswold et al., 2002) acceleration factor, 2; multiband factor, 4; flip angle, 59°; slice matrix size, 106 × 106; voxel size, 2.0 mm isotropic; FOV, 212 × 212 mm; bandwidth: 1745 Hz/px; echo spacing: 68 ms]. To allow for correction of distortions due to magnetic field inhomogeneity, we acquired field maps using a dual echo 2D gradient-echo sequence (64 axial slices; TR, 1020 ms; TE, 10 ms and 12.46 ms; flip angle, 90°; slice matrix size, 64 × 64, slice thickness, 2 mm; FOV, 224 × 224mm). A high-resolution structural image (1 mm isotropic) was acquired using a T1-weighted 3D magnetization-prepared rapid gradient-echo sequence (MP-RAGE; TR, 2.3 s; TE, 3.03 ms; flip angle, 8°; FOV, 256 × 256 × 192 mm).

### MRI data preprocessing in standard stereotactic space and group analyses of the fear conditioning paradigm

To investigate BOLD response activation patterns across participants for the general task effects in the fear conditioning paradigm, MRI data were pre-processed in standard stereotactic (MNI152) space (using SPM12; http://www.fil.ion.ucl.ac.uk/spm; Wellcome Department of Imaging Neuroscience, London, UK). For this, structural images were segmented into grey matter, white matter, and CSF images using a unified probabilistic template registration and tissue classification method (Ashburner & Friston, 2005). Tissue images were then registered with site-specific tissue templates (created from 384 T1-weighted scans) using DARTEL (Ashburner, 2007), and registered (using an affine transformation) with the MNI152 template included in SPM12. Identical transformations were applied to all functional images, which were resliced into 2 mm isotropic voxels and smoothed with a 6 mm FWHM Gaussian kernel. For three participants, the MRI data could not be analyzed due to a reconstruction error. These participants were therefore excluded from this analysis.

The first-level model included four regressors of interest (CS+_threat_, CS−_threat_, CS+_safe_, CS−_safe_) for both runs using 5 s box car functions. Additionally, the period during which participants were navigating was modeled in a separate regressor in each run using a box car function with a duration set to the navigation duration of that period. Responses to shocks were modeled in an additional regressor using a delta function. The model additionally included six movement parameter regressors (3 translations, 3 rotations) derived from rigid-body motion correction, 26 RETROICOR physiological noise regressors (see above), high-pass filtering (1/128 Hz cut-off), and AR(1) serial correlations correction. Single-subject contrast maps obtained from first-level analyses were entered into second-level random effects analyses (one-sample t-test). We used a cluster-forming voxel-level threshold of p< .001 (uncorrected). Alpha was set at .05, whole-brain family-wise error (FWE) corrected at the cluster level using Gaussian Random Field Theory-based methods (Friston et al., 1996).

### MRI data preprocessing in native space and statistical analyses of the context training paradigm

For the classification analysis, BOLD-fMRI data was preprocessed in native space (*i.e*., including the steps described above, but without stereotactic normalization) using SPM12 (http://www.fil.ion.ucl.ac.uk/spm; Wellcome Department of Imaging Neuroscience, London, UK). We created a first-level model using a finite impulse response (FIR) model which included 20 time-bins (TR = 0.909 s) time-locked to trials. Bin number 2 to 8 covered the fixation duration and were used for classification of the context. This first-level model makes no assumptions regarding the HRF shape and yields independent response estimates for all time bins.

We used a leave-one-out procedure to model each of the 48 trials separately, meaning 48 first-level models were created for each participant. In each one trial was modeled separately using one 20-regressor FIR set, and all other trials where collapsed into another 20-regressor FIR set (divided over the two runs). The resulting contrast images were used as data input for the classifier, as explained below.

### Linear support vector machine classification, permutation testing, and region of interest definition

A linear support vector machine (SVM) classifier was used to obtain a linear discriminant function distinguishing between the two buildings (Threat context, Safe context) during the context training paradigm. This was done because during this paradigm no shocks where given. Shock-induced movement and threat-induced arousal could therefore not influence the classification. We took a region-of-interest (ROI) approach based on previous studies showing it is possible to decode environment-specific representantions from the hippocampus (Hassabis et al., 2009). The hippocampus was individually defined in native space using automated anatomical segmentation of T1-weighted images using FreeSurfer 5.3 (http://surfer.nmr.mgh.harvard.edu/). As a control region we took the amygdala, since this region is located close to the hippocampus, but no context decoding can be expected from this region.

First, all features (i.e., voxels within the hippocampus) were scaled using a z-transformation. The mean and standard deviation were always obtained from the training data and applied to both the training and test data. Thus, test data did not influence the scaling. The parameter C, the optimization parameter for misclassifications, was fixed at C=1. We used the train and prediction functions from the LIBSVM toolbox for Matlab to perform the classifications (http://www.csie.ntu.edu.tw/cilin/libsvm/). We trained and tested the data from all trials (24 per building) using a leave-one-out cross-validation procedure with voxels from the predefined hippocampal ROI. We repeated this procedure 1,000 times resulting in an estimate of the average classification accuracy. For statistical testing, we performed a permutation test (1,000 repetitions) for each participant. Before each classification we randomly shuffled the labels. We obtained a standard (z) score for each classification accuracy within the permutation distribution. The resulting z-score was tested against the classification of a control region, namely the amygdala, and against zero.

To assess interindividual differences in context representation and fear generalization, we divided participants into two groups based on a z-score cut-off of 1.645 (95%). Poor context representation was defined as an average (left and right hemisphere) classification z-score of <1.645 (n=14) and a good context representation by an average (left and right hemisphere) classification z-score of >1.645 (n=7).

## Results

### Psychophysiological measures

To confirm the effectiveness of the fear conditioning paradigm, we compared the average skin conductance responses to the CS+ and CS− in both threat and safe contexts using a repeated-measures ANOVA with CS type (CS+ and CS−) and Context (Threat, Safe) as within-subject factors. We found higher average SCR amplitudes for the CS+ compared to the CS− [main effect of CS, *F*(1,20)= 18.36, *p*=3.61E-04, 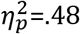]. As expected, these differential responses (CS+ > CS−) were stronger in the threat context compared to the safe context [*F*(1,20)=9.78, *p*=.005, 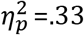]. Nevertheless, differential responses were not only present in the threat context [*t*(20)=3.982, *p*=.001, *D*=1.064], but also in the safe context [*t*(20)=3.447, *p*=.003, *D*=.921]. As an exploratory analysis, we added sex as a between-subject factor, however, there were no significant main effects of, or interactions with sex.

We next reasoned that by averaging across the entire time course of learning, we may not capture fear generalization well, since participants in the beginning have not learned yet which context is safe. Thus, to specifically investigate the relationship between context learning and fear generalization, we used the last two learning trials. We again found higher SCR amplitudes for the CS+ compared to the CS− [main effect of CS, *F*(1,20)= 7.327, *p*=.014, 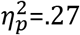]. There was, however, no CS (CS+, CS−) by Context (Threat, Safe) interaction [*F*(1,20)= 1.111, *p*=.31, 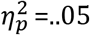]. Analyzing differential SCRs separately within the two contexts, we found a trend-level differential (CS+ > CS−) response in the threat context [*t*(20)=2.030, *p*=.056, *D*=.622] and a significant differential (CS+ > CS−) response in the Safe building [*t*(20)=2.504, *p*=.021, *D*=.773]. Together, this pattern of results indicates that there was robust generalization of conditioned fear expression from the threat context to the safe context by the end of acquisition. For further analyses, we calculated a participant-specific fear-generalization index from the last two trials of acquisition [Δ SCR safe minus Δ SCR threat]. This means that the higher this index was, the stronger the expression of differential conditioned fear responses were in the safe context, relative to the threat context.

In sum, electrodermal data show that our virtual reality fear conditioning paradigm elicited robust conditioned fear responses, and that fear responses generalized to a safe context. ***See Figure 2***.

**Figure 2.**
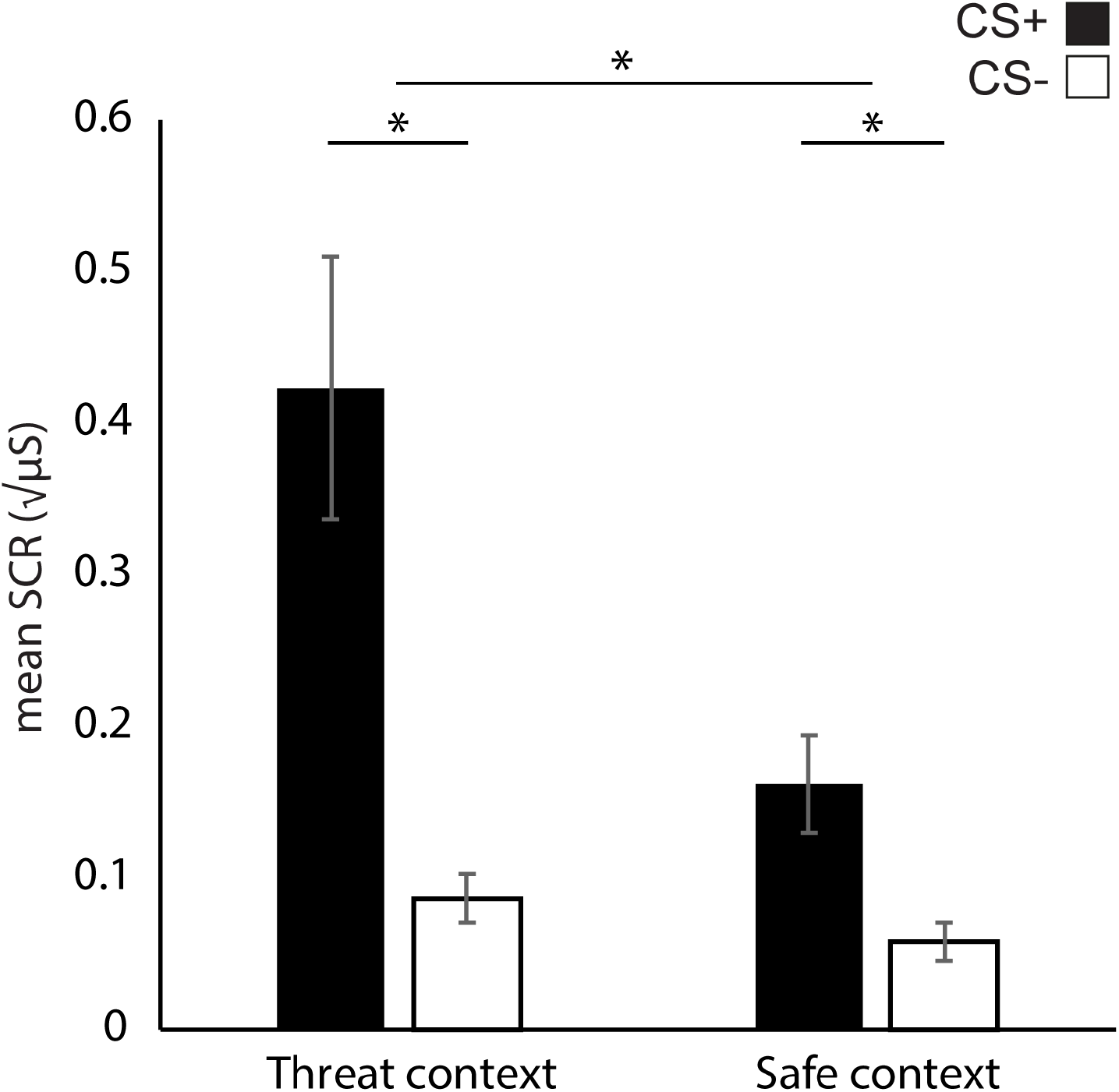
Skin conductance responses (SCR) to CS+ and CS− stimuli, measured in both the threat and safe context. Error bars represent ± standard error of the mean. *= p < .05.

### Univariate functional MRI analysis

We verified whether the fear conditioning paradigm within the virtual reality environment exhibited the expected task-related activity during CS presentation using conventional group analyses in standard stereotactic (MNI152) space. With a whole-brain analysis we first identified regions that were more responsive to the CS+ versus the CS−. In line with results commonly seen in fear conditioning paradigms, we observed robust differential BOLD responses in the left and right anterior insula, and dorsal anterior cingulate cortex [cluster size=125784 mm3, cluster *p*<.001, whole-brain FWE-corrected] as well as the left [cluster size=72 mm3, cluster *p*=.016, FWE-SVC] and right amygdala [cluster size=88 mm3, cluster *p*=.015, FWE-SVC], among others. For the reversed contrast (CS− > CS+) we found differential BOLD responses in the ventromedial prefrontal cortex [cluster size=2688 mm3, cluster *p*<.001, whole-brain FWE-corrected), and left [cluster size= 2464mm3, cluster *p*<.001, whole-brain FWE-corrected] and right angular gyrus [cluster size= 2424 mm3, cluster *p*<.001, whole-brain FWE-corrected], among others. ***See Figure 3 and Table 1***.

**Figure 3.**
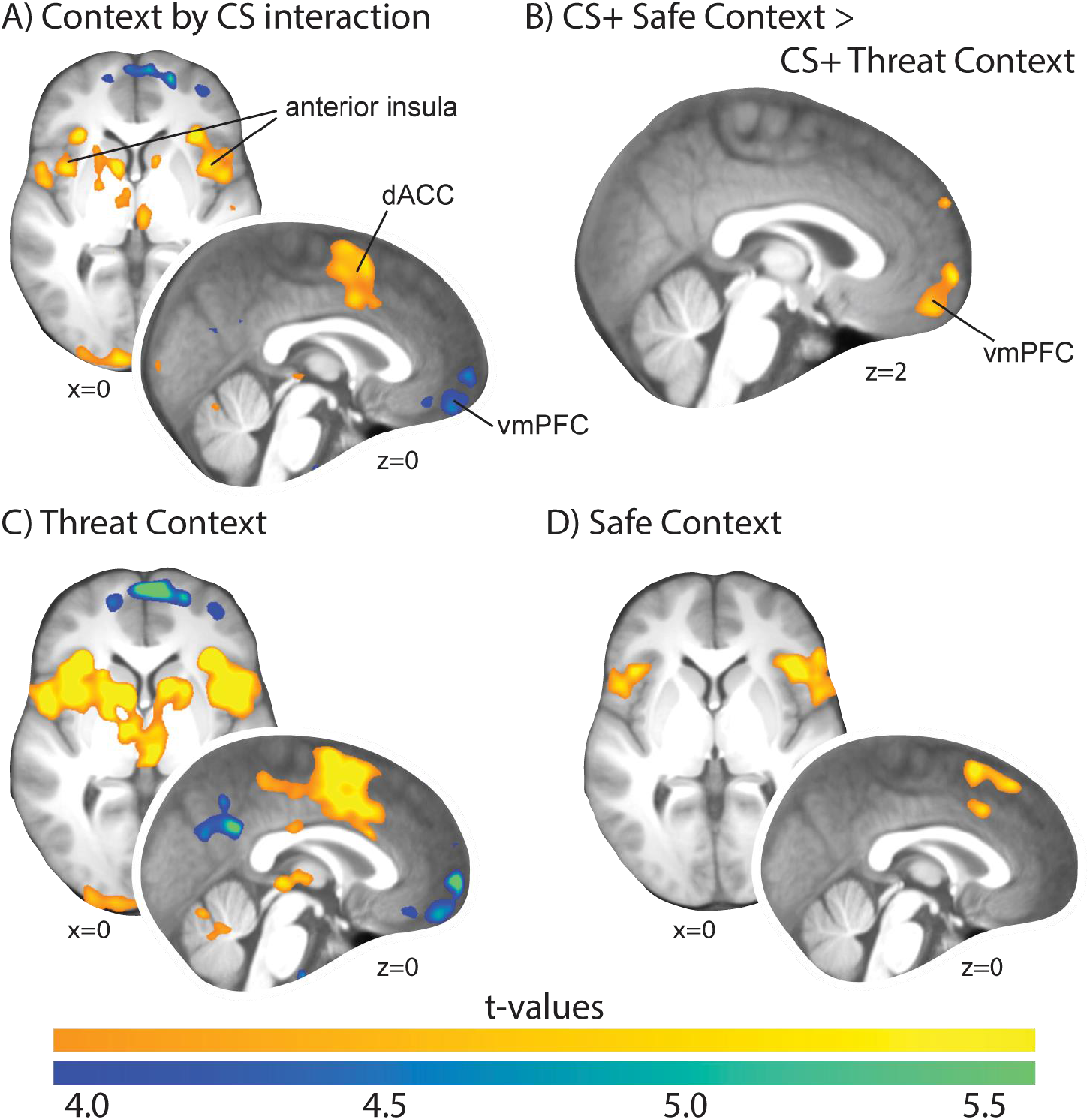
Group statistical parametric maps of differential fear responses. (**A**) Significant clusters in the dorsal anterior cingulate cortex, anterior insula, and thalamus (among others) responded stronger to the CS+ compared to the CS− in the threat context compared to the safe context and ventral medial prefrontal cortex in the revered contrast. (**B**) Significant ventromedial prefrontal cortex cluster in responses the CS+ in the safe compared to the threat context. (C) Significant differential responses (CS+ versus CS−) in the threat context. (D) Significant differential responses (CS+ versus CS−) in the safe context. Statistical parametric maps are thresholded at p < .001, uncorrected, for visualization purposes. Whole-brain cluster-level corrected inferential statistics are reported in Table 1. Maps are overlaid onto the averaged normalized T1-weighted image of all participants.

**Table 1.**
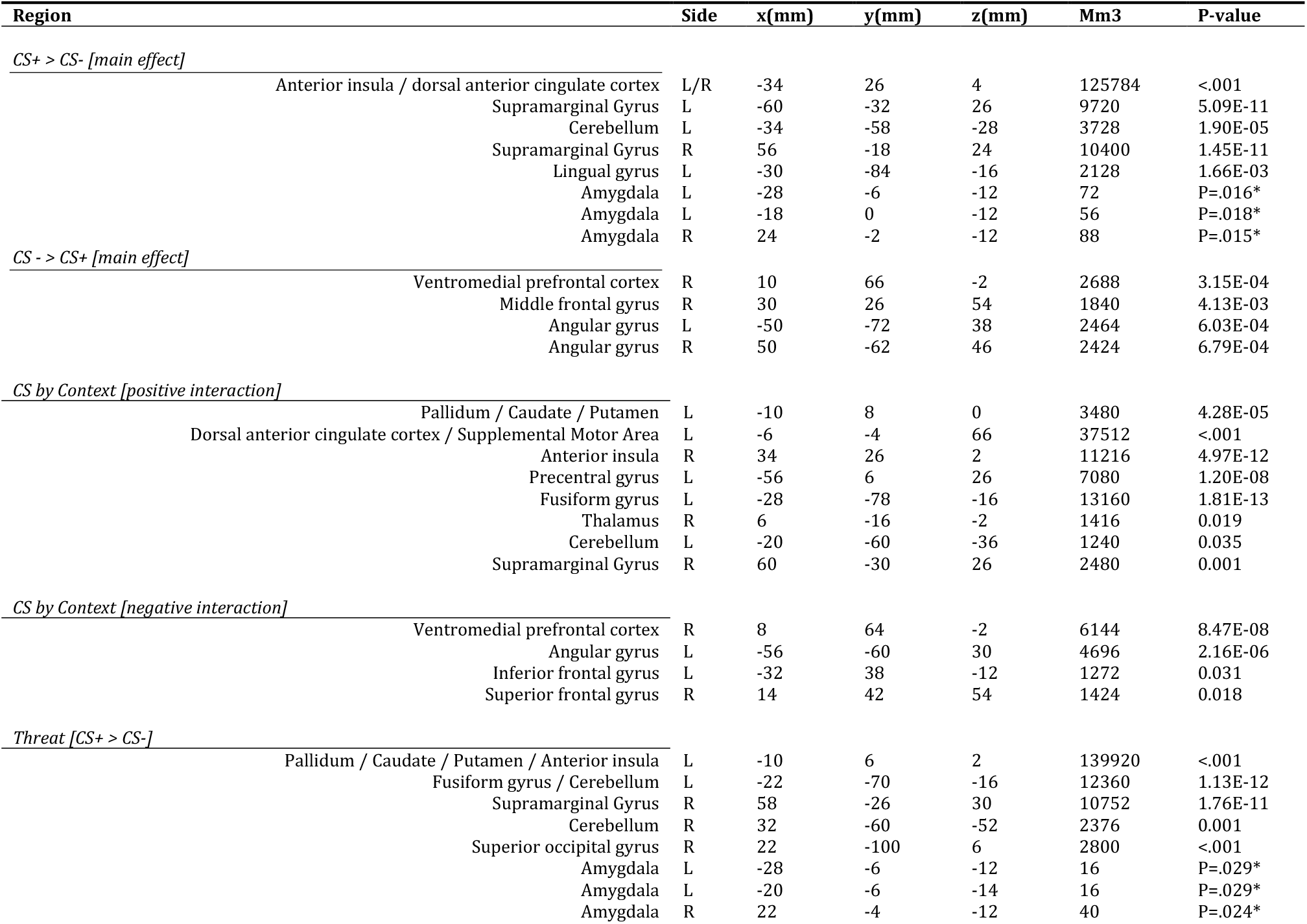
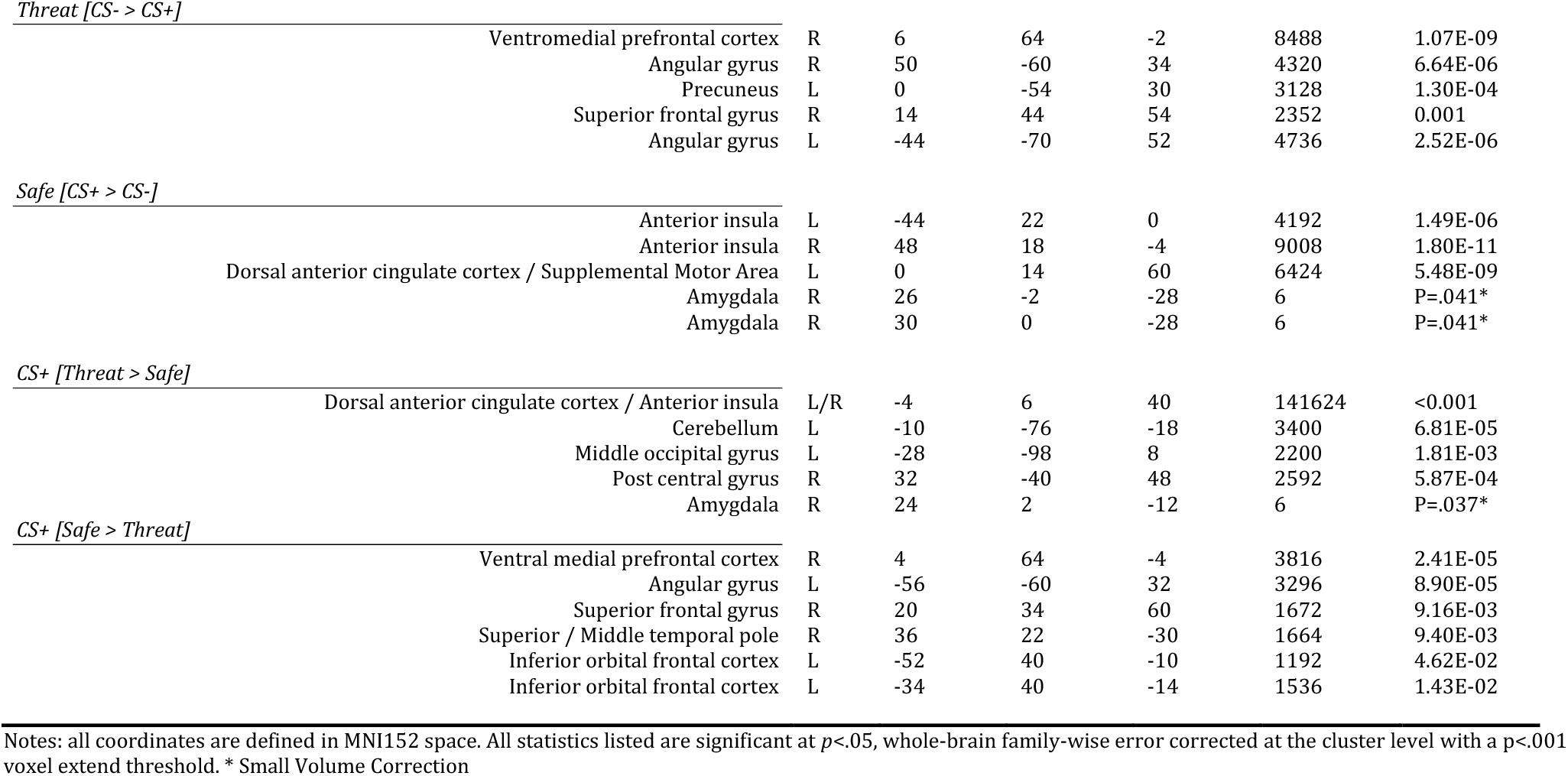
Peak voxel coordinates and cluster statistics of the virtual reality fear conditioning paradigm

| Region | Side | x(mm) | y(mm) | z(mm) | Mm3 | P-value |
| --- | --- | --- | --- | --- | --- | --- |
| *CS+ > CS- [main effect]* |  |  |  |  |  |  |
| Anterior insula / dorsal anterior cingulate cortex | L/R | -34 | 26 | 4 | 125784 | <.001 |
| Supramarginal Gyrus | L | -60 | -32 | 26 | 9720 | 5.09E-11 |
| Cerebellum | L | -34 | -58 | -28 | 3728 | 1.90E-05 |
| Supramarginal Gyrus | R | 56 | -18 | 24 | 10400 | 1.45E-11 |
| Lingual gyrus | L | -30 | -84 | -16 | 2128 | 1.66E-03 |
| Amygdala | L | -28 | -6 | -12 | 72 | P=.016* |
| Amygdala | L | -18 | 0 | -12 | 56 | P=.018* |
| Amygdala | R | 24 | -2 | -12 | 88 | P=.015* |
| *CS- > CS+ [main effect]* |  |  |  |  |  |  |
| Ventromedial prefrontal cortex | R | 10 | 66 | -2 | 2688 | 3.15E-04 |
| Middle frontal gyrus | R | 30 | 26 | 54 | 1840 | 4.13E-03 |
| Angular gyrus | L | -50 | -72 | 38 | 2464 | 6.03E-04 |
| Angular gyrus | R | 50 | -62 | 46 | 2424 | 6.79E-04 |
| *CS by Context [positive interaction]* |  |  |  |  |  |  |
| Pallidum / Caudate / Putamen | L | -10 | 8 | 0 | 3480 | 4.28E-05 |
| Dorsal anterior cingulate cortex / Supplemental Motor Area | L | -6 | -4 | 66 | 37512 | <.001 |
| Anterior insula | R | 34 | 26 | 2 | 11216 | 4.97E-12 |
| Precentral gyrus | L | -56 | 6 | 26 | 7080 | 1.20E-08 |
| Fusiform gyrus | L | -28 | -78 | -16 | 13160 | 1.81E-13 |
| Thalamus | R | 6 | -16 | -2 | 1416 | 0.019 |
| Cerebellum | L | -20 | -60 | -36 | 1240 | 0.035 |
| Supramarginal Gyrus | R | 60 | -30 | 26 | 2480 | 0.001 |
| *CS by Context [negative interaction]* |  |  |  |  |  |  |
| Ventromedial prefrontal cortex | R | 8 | 64 | -2 | 6144 | 8.47E-08 |
| Angular gyrus | L | -56 | -60 | 30 | 4696 | 2.16E-06 |
| Inferior frontal gyrus | L | -32 | 38 | -12 | 1272 | 0.031 |
| Superior frontal gyrus | R | 14 | 42 | 54 | 1424 | 0.018 |
| *Threat [CS+ > CS-]* |  |  |  |  |  |  |
| Pallidum / Caudate / Putamen / Anterior insula | L | -10 | 6 | 2 | 139920 | <.001 |
| Fusiform gyrus / Cerebellum | L | -22 | -70 | -16 | 12360 | 1.13E-12 |
| Supramarginal Gyrus | R | 58 | -26 | 30 | 10752 | 1.76E-11 |
| Cerebellum | R | 32 | -60 | -52 | 2376 | 0.001 |
| Superior occipital gyrus | R | 22 | -100 | 6 | 2800 | <.001 |
| Amygdala | L | -28 | -6 | -12 | 16 | P=.029* |
| Amygdala | L | -20 | -6 | -14 | 16 | P=.029* |
| Amygdala | R | 22 | -4 | -12 | 40 | P=.024* |
| *Threat [CS- > CS+]* |  |  |  |  |  |  |
| Ventromedial prefrontal cortex | R | 6 | 64 | -2 | 8488 | 1.07E-09 |
| Angular gyrus | R | 50 | -60 | 34 | 4320 | 6.64E-06 |
| Precuneus | L | 0 | -54 | 30 | 3128 | 1.30E-04 |
| Superior frontal gyrus | R | 14 | 44 | 54 | 2352 | 0.001 |
| Angular gyrus | L | -44 | -70 | 52 | 4736 | 2.52E-06 |
| *Safe [CS+ > CS-]* |  |  |  |  |  |  |
| Anterior insula | L | -44 | 22 | 0 | 4192 | 1.49E-06 |
| Anterior insula | R | 48 | 18 | -4 | 9008 | 1.80E-11 |
| Dorsal anterior cingulate cortex / Supplemental Motor Area | L | 0 | 14 | 60 | 6424 | 5.48E-09 |
| Amygdala | R | 26 | -2 | -28 | 6 | P=.041* |
| Amygdala | R | 30 | 0 | -28 | 6 | P=.041* |
| *CS+ [Threat > Safe]* |  |  |  |  |  |  |
| Dorsal anterior cingulate cortex / Anterior insula | L/R | -4 | 6 | 40 | 141624 | <0.001 |
| Cerebellum | L | -10 | -76 | -18 | 3400 | 6.81E-05 |
| Middle occipital gyrus | L | -28 | -98 | 8 | 2200 | 1.81E-03 |
| Post central gyrus | R | 32 | -40 | 48 | 2592 | 5.87E-04 |
| Amygdala | R | 24 | 2 | -12 | 6 | P=.037* |
| *CS+ [Safe > Threat]* |  |  |  |  |  |  |
| Ventral medial prefrontal cortex | R | 4 | 64 | -4 | 3816 | 2.41E-05 |
| Angular gyrus | L | -56 | -60 | 32 | 3296 | 8.90E-05 |
| Superior frontal gyrus | R | 20 | 34 | 60 | 1672 | 9.16E-03 |
| Superior / Middle temporal pole | R | 36 | 22 | -30 | 1664 | 9.40E-03 |
| Inferior orbital frontal cortex | L | -52 | 40 | -10 | 1192 | 4.62E-02 |
| Inferior orbital frontal cortex | L | -34 | 40 | -14 | 1536 | 1.43E-02 |

Notes: all coordinates are defined in MNI152 space. All statistics listed are significant at $p<.05$ , whole-brain family-wise error corrected at the cluster level with a $p<.001$ voxel extend threshold. \* Small Volume Correction

Additionally, we found an interaction with Context (Threat context, Safe context) in similar regions, including the dorsal anterior cingulate cortex [cluster size=37512 mm3, cluster *p*<.001, whole-brain FWE-corrected] as well as the ventromedial prefrontal cortex [cluster size=6144 mm3, cluster *p*<.001, whole-brain FWE-corrected], among others. When investigating the differential responses (CS+ versus CS−) in both contexts separately, we found differential responses in the threat context in the anterior insula, dorsal anterior cingulate cortex, caudate and putamen [cluster size=139920 mm3, cluster *p*<.001, whole-brain FWE-corrected], left [cluster size=16 mm3, cluster *p*=.029, FWE-SVC] and right amygdala [cluster size=40 mm3, cluster *p*=.024, FWE-SVC], as well as the vmPFC [cluster size=8488 mm3, cluster *p*<.001, whole-brain FWE-corrected], among others. ***See Figure 3 and Table 1***.

Critically, we also observed differential responses within the safe context (i.e., where participants never received a shock) in the left [cluster size=4192 mm3, cluster *p*<.001, whole-brain FWE-corrected] and right anterior insula [cluster size=9008 mm3, cluster *p*<.001, whole-brain FWE-corrected], and left [cluster size=6 mm3, cluster *p*=.041, FWE-SVC] and right amygdala [cluster size=6 mm3, cluster *p*=.041, FWE-SVC], among others. Finally, as predicted, when comparing the CS+ in the safe building with the CS+ in the threat building, we saw increased responses in the ventromedial prefrontal cortex [cluster size=3816 mm3, cluster *p*<.001, whole-brain FWE-corrected]. ***See Figure 3 and Table 1***.

In sum, similar to the SCR data, our virtual reality paradigm elicited robust fear conditioning responses as well as generalized fear responses in the safe context. Moreover, the CS+ in the safe context elicited increased activation in the vmPFC, a region critically implicated in fear extinction.

### Context classification

Finally, to test our hypothesis regarding the association between strength of context representations and fear generalization, we trained and tested a linear support vector machine (SVM) on voxels from the hippocampus in the context training paradigm, before participants received any shock. We found an average (1000 repetitions of this procedure) of 54.20% accuracy (left: 54.31 % and right: 54.09 %). For statistical testing, we repeated the procedure but this time before each classification we shuffled the labels, resulting in a permutation distribution for each individual (1000 repetitions). Next, we obtained a standard score [(accuracy – average permutation accuracy) / standard deviation permutation] for each individual and each hemisphere. This resulted in an average z-score of 1.33 (left: 1.37 and right: 1.29). As an additional control analysis, we repeated the entire procedure for a control region. We took the amygdala since this region is bordering the hippocampus, is involved in fear learning, but not involved in context representation. We found an average accuracy of 52.00% (left: 52.21 and right: 51.80) and average z-score of 0.55 (left: 0.61 and right: 0.48). We then performed a ROI (hippocampus, control) by Hemisphere (left, right) repeated-measures ANOVA and found that z-scores were higher in the hippocampus than the control region [*F*(1,20)=43.51, *p*<.001, *P*η^2^=.68]. Follow-up one-sample t-test (corrected for four comparisons) showed that for the hippocampus, z-scores were significantly above chance [left: *t*(20)=6.80=, *p*<.001, corrected, and right: *t*(20)=5.774, *p*<.001, corrected], which was not the case for the control region [left: *t*(20)=2.388, *p*=.11, corrected, and right: *t*(20)=2.052, *p*=.20, corrected]. ***See Figure 4D***.

**Figure 4.**
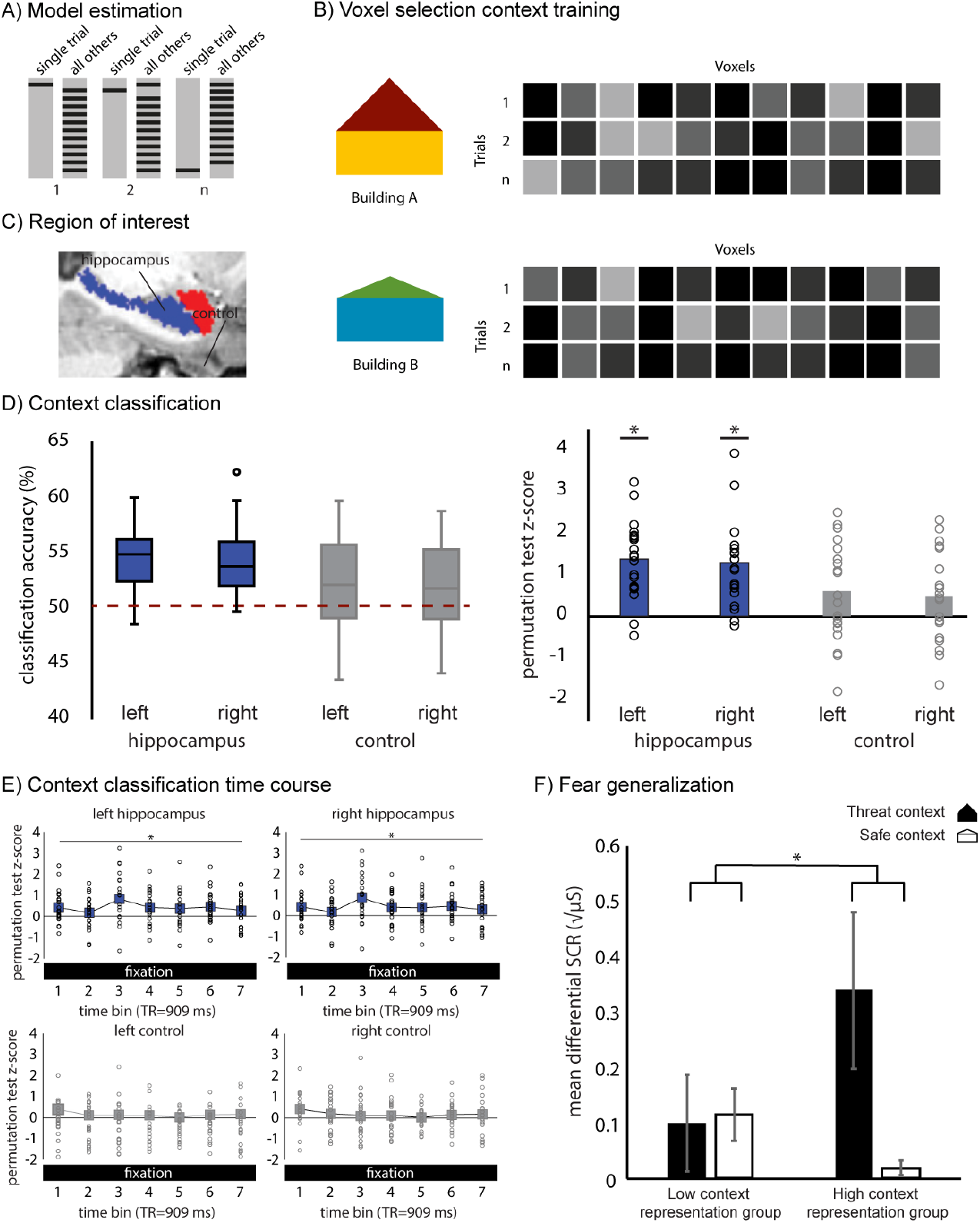
Classification procedure. (**A**) During the context training paradigm, each trial was modeled in a separate GLM. Next, voxels from the hippocampus (and amygdala as a control region) were extracted from each contrast image for each trial and each bin. A linear SVM was trained and tested using a leave-one-out cross-validation procedure. (**B**) Schematic overview of the voxel selection. (**C**) Anatomical individual segmentation of the hippocampus using FSL first and the amygdala as a control region. (**D**) Classification accuracy between the two buildings for voxels in the hippocampus (and amygdala as control). (**E**) Classification accuracy (z-score) per time bin. (**F**) Differential skin conductance responses (SCR) of the last two trials measured in both the threat and safe context separated for the low (n=14) and high (n=7) context representation groups. Error bars represent ± standard error of the mean. *= p < .05 using permutation testing.

Finally, we investigated the time course of BOLD signal within trials by performing the same classification procedure on each time bin within the 7-s fixation period for the hippocampus and the control region. We performed an ROI (hippocampus, control) by Hemisphere (left, right) by Time bin (7 bins) repeated-measures ANOVA and found that z-scores were higher in the hippocampus than the control region [*F*(1,20)=15.50, *p*<.001, *P*η^2^=.44]. Follow-up tests (corrected for two comparisons) showed z-scores were significantly above zero for the hippocampus [*F*(1,20)=14.24, *P*=.002, corrected, *p*η^2^=.42], which was not the case for the amygdala [*F*(1,20)=2.311, *P*=.29, corrected, *p*η^2^=.10]. ***See Figure 4E***.

In sum, in line with the previous literature, we found distinguishable multi-voxel patterns of hippocampal environment-specific representantions within a virtual reality environment.

### Context representations associated with fear generalization: SCR

Finally, we reasoned that a low context representations would predict poor fear contextualization. To investigate this, we made use of the fear-generalization index described above, which was based on generalization of conditioned fear to the safe context in the final two trials of acquisition. To determine to what extent participants exhibited strong versus weak context representations, we divided participants into two groups based on a z-score cut-off of 1.645 (95%) for the individual classifier performance in distinguishing the two contexts. In this way we defined a group of low context representation (z-score < 1.645, n=14) and a group of high context representation (z-score> 1.645, n=7). Next, we added Context representations as a between-subject factor to the SCR analyses (as described above).

We found stronger fear generalization in the low context representation group compared to the high context representation group [*t*(19)=1.806, *p*=.04, one-tailed, D=.836]. Due to a violation of the normality assumption, we performed a permutation test and confirmed this result [*t*= 1.717, *p*=.04, one-tailed permutation test, 100,000 repetitions]. Further testing showed that this effect was not specific to either of the two contexts, meaning this effect may be partly driven by numerically stronger differential SCRs in the safe context for the low context representation group, but also partly by numerically stronger differential SCRs in the threat context in the high context representation group. ***See Figure 4F***.

Additionally, we tested whether fear acquisition in general was affected by context representation. We found a CS (CS+, CS−) by Context (Threat context, Safe context) by Context representation (Low, High) interaction [*F*(1,19)=5.391, *p*=.032, 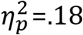] across all learning trials. Differential responses (CS+ minus CS−) in the Threat context was stronger for the high context representation group compared to the low context representation group [*F*(1,19)=5.084, *p*=.036, 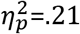], but there were no differences in differential responses (CS+ minus CS−) in the Safe context [*F*(1,19)=.315, *p*=.58, 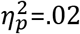] between the groups. This suggests that contextual information also plays a role during fear acquisition.

In conclusion, we found that the high context representation group showed stronger differentially conditioned SCRs in a threat context overall, indicating more strongly discriminative fear learning. We furthermore found evidence that the low context representation group showed stronger differential responses in a safe context, reflecting stronger fear generalization.

## Discussion

This study was designed to investigate the role of spatial context representations in context-dependent expression and generalization of learned fear. We hypothesized that suppression of the expression of conditioned fear in safe contexts would depend on similar neural mechanisms as those involved in extinction recall. Furthermore, we hypothesized that a low context representation of a virtual environment would underlie the generalization of fear to a safe context. We tested these hypotheses in a contextual fear conditioning paradigm in which context in a self-navigated virtual environment served as an “occasion setter”.

We report the following findings. First, we found expression of differentially conditioned responses (*i.e*., increased SCRs to the CS+ compared to the CS−) in a safe context (*i.e*., where participants never received a shock), demonstrating generalization of fear and thus validating our paradigm. Second, in line with our expectation that context-dependent suppression of fear is similar to extinction recall, we found increased ventromedial prefrontal cortex (vmPFC) responses to the CS+ in the safe context compared to the CS+ in the threat context. Third, we found above-chance decoding of environment-specific representations within the virtual environment (measured before fear acquisition took place). Lastly, interindividual differences in context representation predicted later differential fear conditioning strength, independent of the context. Critically, weaker context representations prior to fear learning predicted later fear generalization.

We found increased SCRs as well as increased BOLD responses in regions typically activated in response to cues associated with threat, in a safe context (*i.e*., where no aversive event has taken place). Heightened fear responses in a safe environment is one of the hallmarks of fear and stress-related disorders (Jovanovic, Kazama, Bachevalier, & Davis, 2012). Here, we found that such a response to a potentially threatening cue in a safe context is present in a healthy population. We extend previous findings of fear generalization that have shown generalized fear responses to perceptually similar CSs (Dunsmoor, Mitroff, & LaBar, 2009; Lissek et al., 2008; Struyf, Zaman, Hermans, & Vervliet, 2017), which were additionally shown to be accompanied by increased BOLD responses in regions implicated in fear acquisition (Dunsmoor, Prince, Murty, Kragel, & LaBar, 2011). We also extend a previous study that has shown fear responses to a CS associated with an aversive event in a safe context (Baas et al., 2004), by showing these generalized physiological responses are accompanied by increased activity in neural systems involved in fear learning.

In line with our hypothesis regarding the role of the vmPFC in context-dependent expression of conditioned fear, we observed increased vmPFC responses to the CS+ in the safe context compared to the CS+ in the threat context. The vmPFC is critically implicated in extinction of fear memory, as detailed in animal models (Milad & Quirk, 2012). Human studies using neuroimaging have indicated that vmPFC responses increase over the course of extinction, in response a to CS+ that no longer predicts an aversive outcome (Phelps et al., 2004). Increased vmPFC responses were also found in response to an extinguished CS+ compared to a non-extinguished CS+, implicating a role of the vmPFC in extinction recall (Milad et al. 2007). It is thought that extinction learning does not overwrite the orginal fear memory (*i.e*., US-CS association), but creates a new safety memory that may be context-dependent (Bouton 2004). It has been suggested that the extinction-induced vmPFC responses following extinction are therefore also context-dependent (Kalish et al., 2006). The increased vmPFC responses we found to a cue signaling threat in a safe context suggest that context-dependent safety learning involves a similar neural pathway as extinction learning. We extend previous findings, however, by showing that vmPFC responses do not per se progress over time but can flexibly shift across a threat and safe context. Thus, our findings indicate that the role of the vmPFC extends beyond one-directional extinction and suppression of fear expression, and is better understood as implementing the capacity to flexibly regulate context-dependent expression of fear.

We found above-chance classification of environment-specific represantations within a virtual reality environment from multi-voxel patterns of BOLD-fMRI recorded within the hippocampus. This is in line with previous studies (Hassabis et al. 2009; Kim et al., 2017; Rodriguez, 2010; Sulpizio et al. 2014). One question that often arises is whether neuronal representations of the spatial environment, which are known to be coded across distributed neuronal ensembles within the hippocampus, can truly be picked up using a relatively coarse technique such as BOLD-fMRI. Indeed, it is possible that environment-specific represantations could be decoded based on other features than a purely spatial code (e.g., multimodal associations reinstated by exposure to a context). Notably, however, it was recently found that in addition to sinlge-neuron firing rates, local-field potentials (LFP) can also be used to decode spatial location in rodents, and this measure appears as reliabe as neural spikes (Agarwal et al 2014). Because LFPs are also thought to underlie the BOLD signal (Magri, Schridde, Murayama, Panzeri, & Logothetis, 2012), it appears reasonable to assume that similar spatial representations could be detected using BOLD as well. Indeed, human neuroimaging studies using BOLD-fMRI in humans have revealed that similar mechanisms are involved in coding space and location as in rodents (Doeller, Barry, Burgess 2010). Namely, using virtual reality paradigms, it was shown that BOLD signal in MTL regions follows a grid-like pattern for space (Doeller, Barry, Burgess 2010), but also in relation to mental stimulation (Bellmund et al., 2017) and visual space (Nau et al., 2018).

Our finding is in line with previous studies (Hassabis et al. 2009; Kim et al., 2017; Rodriguez, 2010; Sulpizio et al. 2014), however, there are a few differences between our design and those of previous studies. First, in our design, participants were able to move freely within the virtual reality environment, and within the context training paradigm, trials start at random locations within the environment to promote path formation (Cei et al., 2014). Second, we have performed statistical testing on z-scores, based on individual permutation distrubutions, and not the individual classification accuracies, because this was shown to be more valid (Haynes, 2015). Third, in our design, we fully controlled for visual confounds. The two buildings were fully identical on the inside, and as input for the classifier we used a seven-second fixation period during which only a fixation dot was visible on the screen.

Finally, we hypothesized that low context representations would be associated with more fear generalization. We indeed found evidence for this association, however, we also found that participants for whom the classifier was able to classify in which building they were in the virtual environment during the training paradigm (*i.e*., before fear acquisition took place) had stronger differential fear responses in the threat context across all learning trials. These two findings are indications of how contextual information can play a dual role in fear acquisition. We propose that, on the one hand, contextual information enhancee adaptive responses to threat by increasing its specificity. Impaired discrimination between threat and safe cues has been linked to high anxiety (Kuhn, Mertens, & Lonsdorf, 2016), suggesting that an increased cue discrimination reflects an adaptive response to threat. Our finding suggests that context representation plays a role in this discrimination. On the other hand, failure to integrate cue and context information into a conjunctive memory may underlie the development of maladaptive threat responses that generalize to safe environments. For instance, the inability to regulate fear using context information is thought to underlie the development of post-traumatic stress disorder (PTSD; Liberzon and Sripada, 2008; Cohen et al., 2009; Acheson et al., 2011). Our findings provide empirical support for this notion by linking poor spatial context representations to the inability to reguate fear responses in a safe context.

In conclusion, we demonstrate generalized conditioned fear responses to a safe context within a virtual reality environment. We found that the vmPFC is activated to threat cues in a safe context, suggesting that this region uses contextual information to flexibly adjust expression of conditioned fear, in a similar fashion as during recall of extinction memory. Furthermore, we observed that good context representations are associated with stronger fear learning in the correct context, while poor context representations were associated with stronger fear generalization (i.e., conditioned fear expression in a safe context). These experimental findings in a healthy population open new avenues for exploring the role of spatial and contextual representations in pathologically generalized fear, as observed for instance in PTSD.

